# Highly Multiplexed Spatial Mapping of Microbial Communities

**DOI:** 10.1101/678672

**Authors:** Hao Shi, Warren Zipfel, Ilana Brito, Iwijn De Vlaminck

## Abstract

Mapping the complex biogeography of microbial communities *in situ* with high taxonomic and spatial resolution poses a major challenge because of the high density and rich diversity of species in environmental microbiomes and the limitations of optical imaging technology. Here, we introduce High Phylogenetic Resolution microbiome mapping by Fluorescence In-Situ Hybridization (HiPR-FISH), a versatile and cost-effective technology that uses binary encoding and spectral imaging and machine learning based decoding to create micron-scale maps of the locations and identities of hundreds of microbial species in complex communities. We demonstrate the ability of 10-bit HiPR-FISH to distinguish 1023 *E. coli* strains, each fluorescently labeled with a unique binary barcode. HiPR-FISH, in conjunction with custom algorithms for automated probe design and segmentation of single-cells in the native context of tissues, reveals the intricate spatial architectures formed by bacteria in the human oral plaque microbiome and disruption of spatial networks in the mouse gut microbiome in response to antibiotic treatment. HiPR-FISH provides a framework for analyzing the spatial organization of microbial communities in tissues and the environment at single cell resolution.

## INTRODUCTION

Microbial communities often exhibit rich taxonomic diversity and exquisite spatial organization (*1*–*4*). While the taxonomic diversity of microbiomes is readily accessible by metagenomic sequencing, the spatial organization of complex microbial communities remains very difficult to survey. Fluorescence in situ hybridization assays that target ribosomal RNA (rRNA) for taxonomic identification and visualization have been developed, but have so far been severely limited in taxonomic resolution. The best in class method, CLASI-FISH, enables detection of up to 15 taxa in environmental samples using hybridization probes tagged with combinations of 6 fluorophores (*5, 6*). These studies highlight the spatial organization of several abundant taxa, but are insufficient for unbiased analysis of microbial biogeography at the microbiome scale in the mammalian gastrointestinal tract, where hundreds to thousands of species coexist (*7*). Refinements in CLASI-FISH enable detection of up to 120 distinctly labeled *E. coli* isolates using 16 different fluorophores but have not been applied to environmental microbial communities and are very costly to scale up (*8*). Studies to-date have furthermore been largely qualitative, with a few quantitative analyses using coarse-grained approaches (*9, 10*), making spatial information at the single cell level inaccessible.

Here, we demonstrate HiPR-FISH, a low-cost microbiome mapping technology that achieves 1000-fold multiplexity in taxa identification and visualization with a single round of imaging on a standard confocal microscope. HiPR-FISH achieves high multiplexity with a binary barcoding scheme to translate taxon identity to n-bit binary words, and a machine learning approach for the classification of the combined spectra of up to 10 fluorophores (Fig. 1A & Supplementary Figures 1 & 2). For binary barcoding of species, HiPR-FISH implements a two-step hybridization scheme, with a first step that uses taxon-specific 16S rRNA probes modified with DNA flanking regions, and a second step with fluorescently labeled readout probes. The high copy number of 16S rRNA in each cell provides flexibility in probe sequence design. The number of flanking sequences per probe sequence is limited to two, and n-bit encoding with n>2 is achieved with a mixture of probe sequences targeting the same rRNA species. Unlike previous approaches that have used species specific probes tagged with fluorophores (*11, 12*), HiPR-FISH uses unmodified probes that can be synthesized with array technologies, leading to a much lower cost per assay design (Fig. 1B).

**Figure 1.**
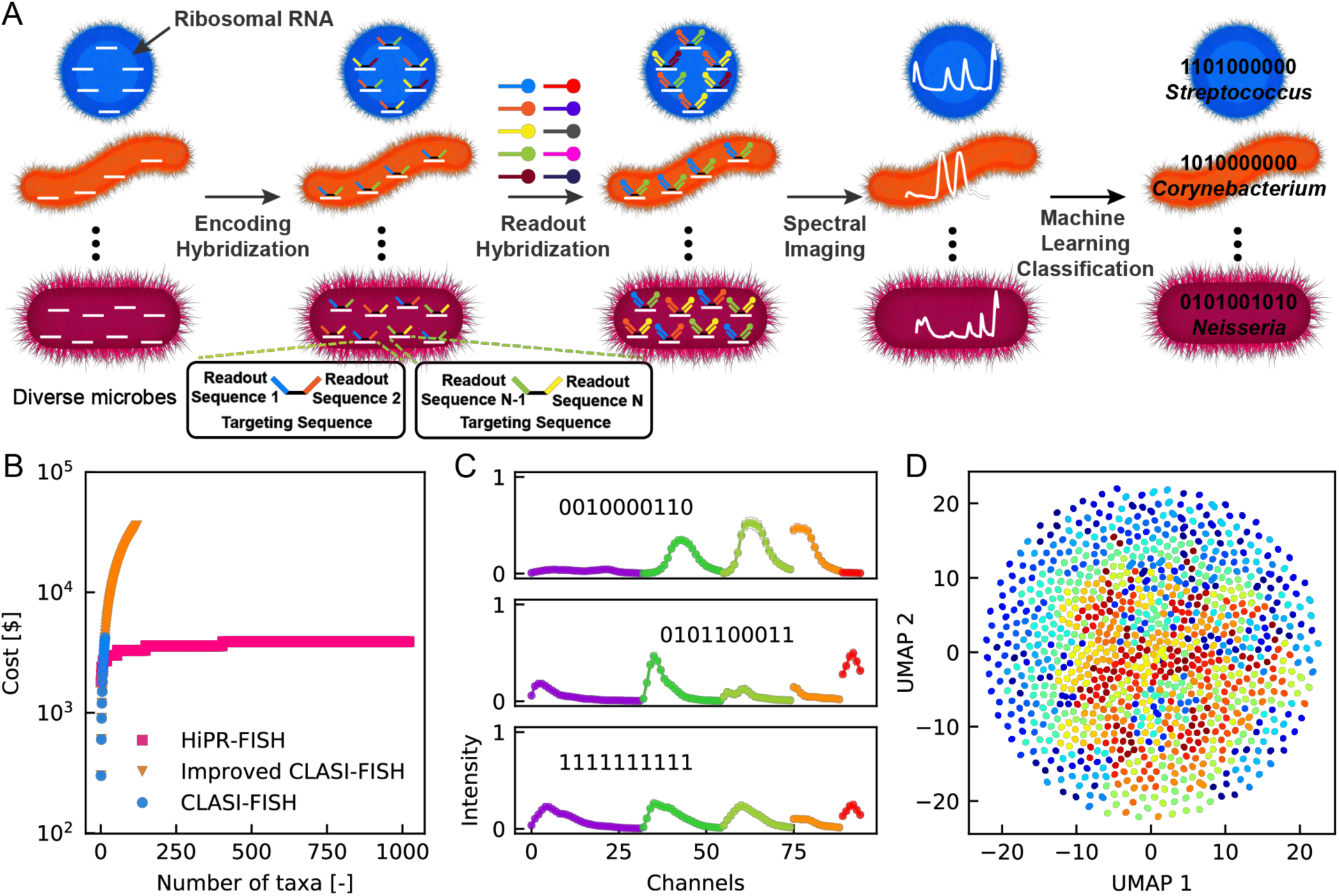
HiPR-FISH working principle. **A.** HIPR-FISH takes advantage of the high abundance of ribosomes in microbial cells to label a given species of bacteria with up to 10 different fluorophores, enabling 1023-plex multiplexity in taxa identification and visualization. **B.** Cost per probe panel as a function of multiplexity for HiPR-FISH compared to other methods. **C.** Five example single cell spectra acquired using spectral imaging. **D.** A UMAP representation of all 1023 experimentally measured spectra using a 10-bit system. Each spectrum is colored sequentially according to their numeric barcode using a jet colormap divided into 1023 segments.

To apply HiPR-FISH to quantitative analyses of environmental microbial communities with single-cell resolution, we developed a routine for automated image segmentation that overcomes the challenges inherent to delineating single cells in microbiomes, including the significant cell-to-cell variability in fluorescence intensity, and the high density and large morphological diversity of microbial cells. The routine, inspired by computer vision approaches to image enhancement, leverages the information contained in directional intensity profiles around a given pixel or voxel to reduce the global dynamic range in fluorescence intensity and enhance edge contrast. Segmentation at the single cell level after HiPR-FISH enabled quantitative physical analysis of microbial species in the human oral plaque microbiome and enabled to unravel single-cell spatial adjacency networks. We used HiPR-FISH to study the spatial architecture of the mouse gut microbiome and found that antibiotics induce marked changes in both abundance and spatial organization.

## RESULTS

### Implementation of HiPR-FISH

For the practical implementation of HiPR-FISH, we used barcodes comprised of up to ten fluorophores with distinct optical excitation and emission spectra, creating up to 1023 unique combinations (2^10^-1, Supplementary Figure 3). We barcoded cells using the two-step hybridization scheme detailed in Figure 1A, and recorded the barcode spectra using a standard confocal microscope. For each cell, we concatenated spectra measured sequentially using five excitation lasers, averaged the spectra measured across all pixels within a cell (Fig. 1C), and decoded the cell barcode using a custom machine-learning classifier. For classification, we first used Uniform Manifold Approximation and Projection (UMAP) to project spectra onto two dimensions (Supplementary Figure 3). A custom excitation-channel-wise cosine distance kernel in UMAP achieved clean separation of all barcodes (Fig. 1D). A support vector machine (SVM) trained on the two-dimensional representation of pure reference spectra was then used for *de novo* barcode prediction. To account for variable ribosomal density in cells from different taxa in more complex systems such as dense environmental biofilms, we use physical models that model Förster Resonance Energy Transfer to refine training spectra. To generate training data for the classifier, we measured the emission spectra for each barcode individually. For HiPR-FISH with 10 fluorophores or bits, this required the generation of 1023 *E. coli* isolates barcoded with 1023 different barcodes.

### Proof-of-principle

To test the robustness of HiPR-FISH, we characterized predefined mixtures of the 1023 *E. coli* barcode strains. We first created and imaged a mixture with all barcoded strains present at equal concentration. We imaged 50 135 μm×135 μm fields of view, comprising a total of 71,465 cells (Figs. 2A-B). We used the above described machine-learning classifier to determine the barcode among the collection of 1023 possible barcodes that best corresponded to the measured spectrum for each cell. We found that all barcodes were represented in the mixture, and the mean fractional abundance of each barcode was ∽0.001, as expected (Fig. 2C). We next randomly divided the 1023 aliquots into 8 groups, where each group comprised 127 or 128 barcodes, and mixed barcodes in the same group at varying relative abundance, ranging from 0.1×10^−3^ to 15.5×10^−3^. We measured 35,000 to 40,000 single cell spectra across 50 fields of view for each group and quantified the relative abundance of different barcodes in each mixture. We again applied the machine learning classifier to determine the barcode that most likely corresponded to each measured single-cell spectrum. We found close agreement between the expected abundance and the measured abundance across all barcodes, with an average slope of 0.94 and an average R^2^ value of 0.87 (Fig. 2D). To quantify the accuracy of the barcode classification routine, we defined the gross error rate as the fraction of all detected barcodes that does not belong in a group of barcodes. We found that barcode mis-assignment is rare, with gross error rates ranging from 0.8% to 6.7%. These findings support the robustness of 10-bit HiPR-FISH.

**Figure 2.**
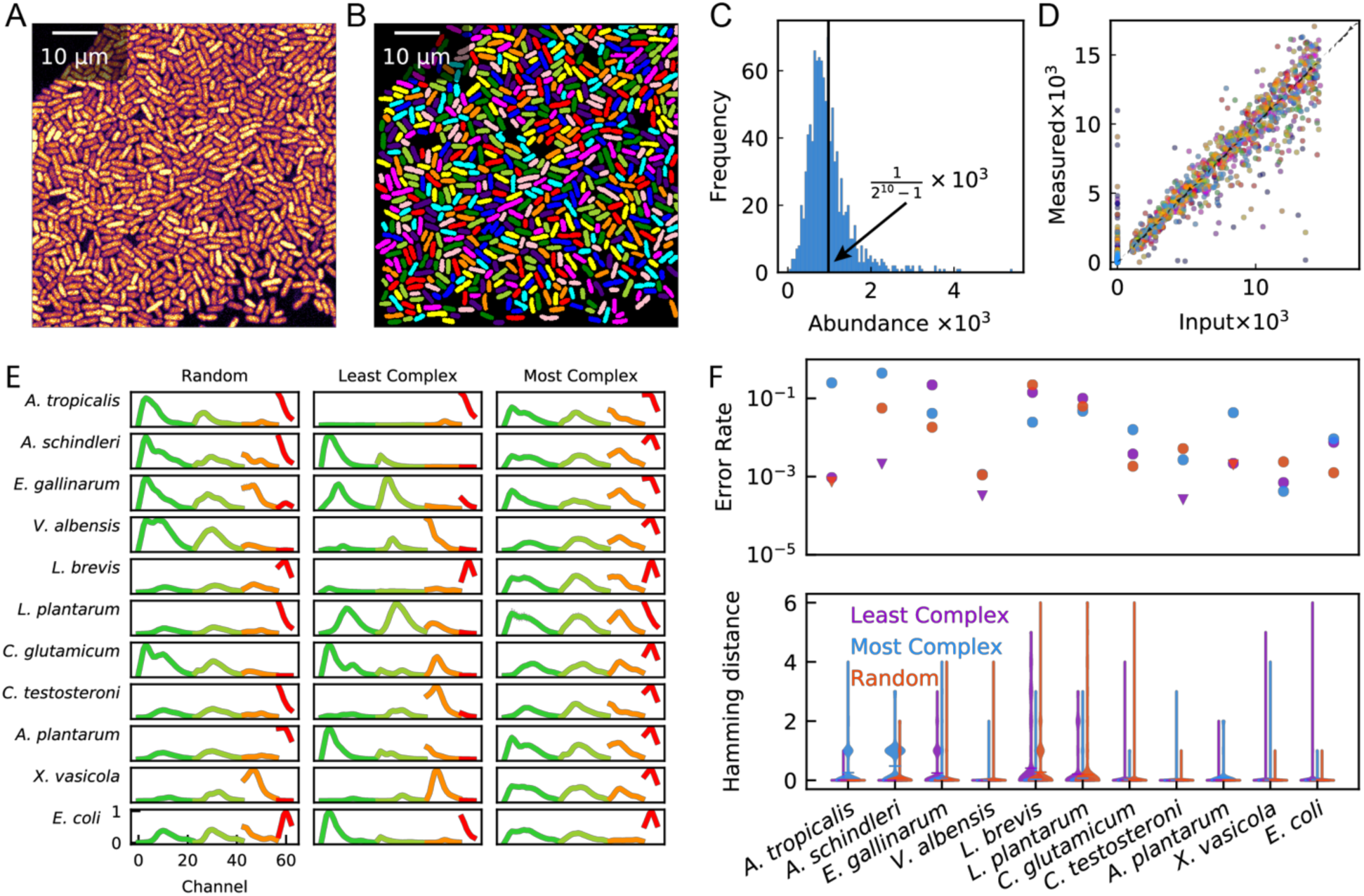
HiPR-FISH proof-of-principle with mock *E. coli* and multi-species communities. **A.** A typical field of view of *E. coli* cells. The raw image has been averaged over the spectral dimension to generate a 2D representation. **B.** Single cell segmentation of the field of view in (A). **C.** Histogram of the relative abundance of an equal-concentration mixture of all 1023 species of barcoded *E. coli* cells. **D.** Correlation of measured abundance and input abundance in *E. coli* synthetic communities. Each community consists of 127 to 128 species of barcoded *E. coli* cells at varying concentrations. **E.** Average spectrum of each of the 11 species in the synthetic community. **F.** Mis-identification rate for each species in the synthetic community. Triangular data points indicate upper limits in cases where there was no mis-identified cell. Hamming distance shows that most incorrect classifications are only one bit away from the correct barcode.

To further demonstrate the principle of HiPR-FISH, we probed and imaged a mock community consisting of 11 species of bacteria. To assess whether HiPR-FISH is compatible with complex environmental samples, where microbes with diverse cell envelope structures coexist, we included both gram positive (n = 4) and gram negative (n = 7) species in this panel. We generated three sets of probes (A, B, C). Each probe set comprised a common list of targeting sequence that are specific to the 11 species in the synthetic community but had different flanking encoding sequences. In particular, targeting sequences in set A, B, and C are encoded with the least complex barcodes, the most complex barcodes, and a random selection of barcodes, respectively, where the complexity of a binary sequence is defined as the number of true bits of that barcode. Barcodes with more true bits are more complex, in the sense that the emission spectrum is a combination of a greater number of fluorophores. To evaluate the specificity of each custom-designed species-specific probe, we hybridized each probe set to pure cultures of each species, recorded spectral images for each probe set-species combination, and classified single cell spectra (Fig. 2E). We find strong agreement between the assigned barcode and expected barcode for all probe set-species combination (Fig. 2F). In cases of spectral misclassification, the Hamming distance between the measured barcode and the assigned barcode is mostly 1, indicating that the misclassified barcode is just one bit away from the correct barcode. Overall, we find that we can achieve species-specific detection and flexible binary barcode encoding using HiPR-FISH in a 11-species synthetic community.

### Probe design and synthesis for environmental microbiome imaging

HiPR-FISH requires the design of 100s-1000s of unique hybridization probes. To make this problem tractable, we developed and implemented an automated bioinformatic workflow for probe library design (Supplementary Figure 4). To inform the design of FISH probe libraries, we generated custom references of full length rRNA genes for each community of interest by PacBio circular consensus sequencing (SMRT-CCS) (*13*) (see Methods). We grouped 16S sequences for each community by taxon and sequence similarity and generated a consensus sequence for each taxon using usearch (*14*). We designed 18-23 bp candidate probes for each consensus sequence using primer3 (*15*). We evaluated the specificity of the candidate probes by alignment against the 16S sequence reference using blastn and removed probes with low specificity (see Methods). Finally, each probe was expanded on both ends with 3 bp spacers, readout sequences, and primer sequences for probe synthesis (see Methods, Supplementary Figure 5). The hybridization probes were electrochemically synthesized in complex pools on arrays of electrodes. Complex pools of hybridization probes comprised of several thousand unique oligos were PCR amplified and converted to ssDNA for in-situ hybridization, as in Ref. (*16*) (Supplementary Fig. 5).

### Image processing and single cell segmentation

Studies of the biogeography of microbial communities have to date been largely qualitative, with only a few examples of quantitative analyses using coarse-grained approaches (*9, 10*). To extract quantitative information with single cell resolution from HiPR-FISH images, we set out to develop streamlined image analysis procedures that require minimal operator input (Fig. 3, Supplementary Figure 6).

**Figure 3.**
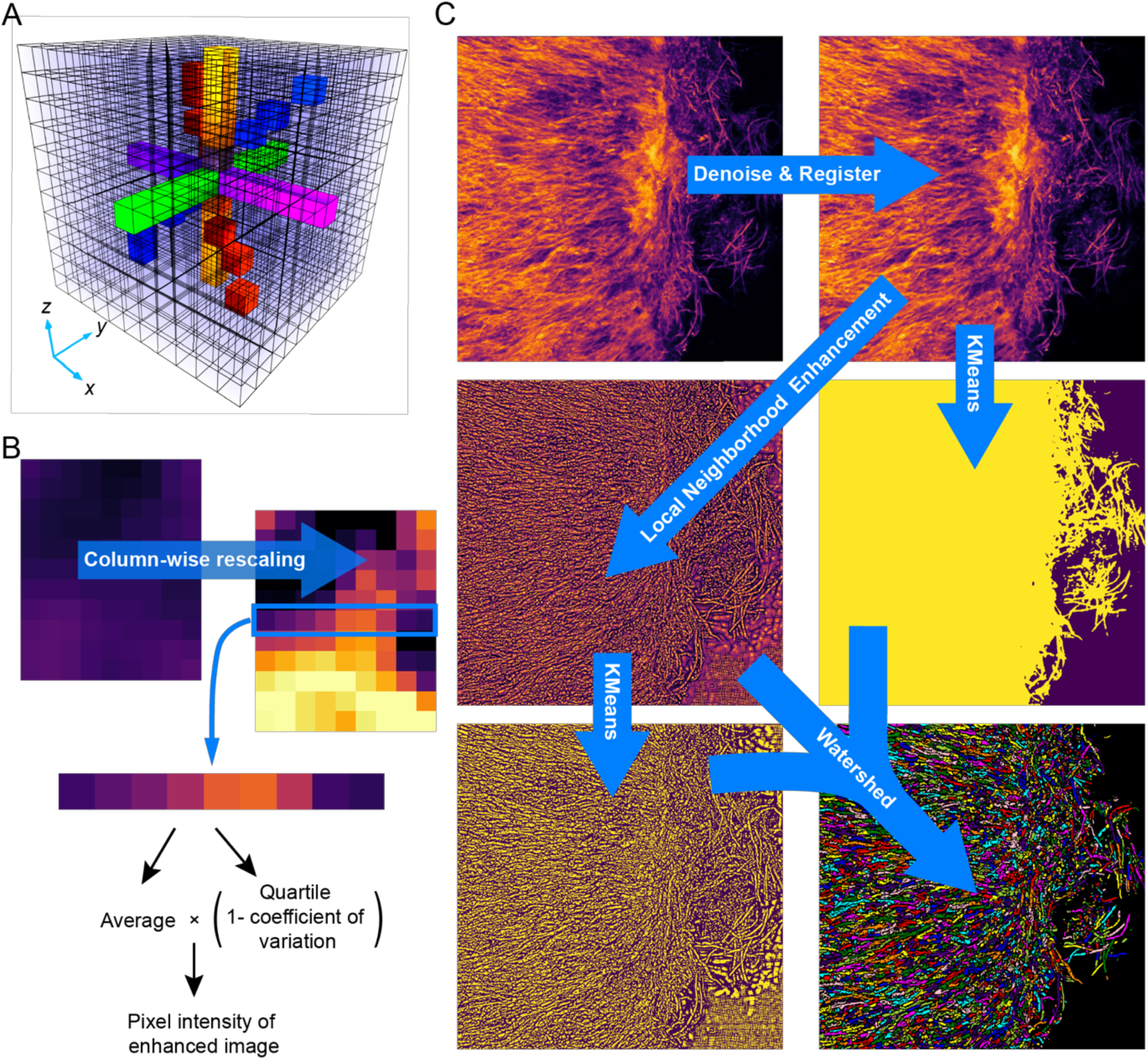
Algorithm for single-cell segmentation. **A.** Example structuring elements used in the local neighborhood enhancement algorithm for 3D images. Voxels of a particular color represent a structuring element centered at the voxel of interest along a specific direction. **B.** An example line profile matrix for a pixel of interest from a 2D biofilm image. Column-wise normalized line profiles enables the calculation of the pixel intensity in the enhanced image. **C.** Key steps in the image processing workflow. Images are denoised using either a time average or a convolutional neural network and registered. Denoised images are enhanced using our custom local neighborhood enhancement algorithm. Background filter masks were generated using the denoised images. Watershed seeds masks were generated using the locally enhanced image. Finally, the enhanced image, the watershed seeds masks, and the background filter masks are used together to segment the denoised image into individual cells.

To identify and segment individual cells, we used the watershed algorithm, which performs well at segmentation tasks, provided that the initial segmentation seed is accurate. The challenge of segmenting densely packed microbes is then reduced to the task of finding a seed image that is closest to the ground truth image. Previous approaches have relied on user-provided parameters and assumptions about morphologies of the objects to be segmented (*17, 18*). We tested the performance of these approaches to create single-cell segmentations of oral biofilms but found that they tend to under-segment communities with diverse shapes, sizes, and ribosome density (Supplementary Figure 7). We therefore developed a new approach to define a seed for watershed segmentation, optimized for HiPR-FISH. This routine attempts to classify each image pixel as belonging to a cell or to the background using information contained in the pixel local neighborhood (Fig. 3). Here, image denoising is first performed using either traditional computer vision approaches such as non-local means (*19*–*21*), or convolutional neural networks (*22, 23*) (for samples where independent replicate measurements are available). Denoised images are registered using phase correlation to reduce blurring due to microscope stage drift (*24*). Next, for each pixel, the intensity line profiles crossing the pixel along multiple directions are extracted and normalized (parameterized by the azimuthal angle ϕ and polar angle θ, Figs. 3A-C). The average and quartile coefficient of variation of the pixel of interest over all normalized intensity line profiles is then calculated to produce the pre-processed image, which exhibits a low global dynamic range and sharp boundaries between closely packed cells. Next, pixels in the enhanced image are classified into two brightness categories using k-means clustering, where the brighter pixels form the seed image for watershed. To exclude background pixels, a filter mask based on the natural log of the raw intensity of the pixels is applied. Finally, the seeds, the enhanced image, and the background filter mask are used to produce a final segmentation using watershed. We found that this algorithm performs much better at single-cell segmentation of densely packed microbes with diverse shapes and ribosome density than previously reported approaches (Supplementary Figure 7).

### Spatial organization in environmental microbiomes

We next applied HiPR-FISH to create spatial maps of the locations and identities of microbial species in human plaque biofilms. The oral cavity hosts one of the most diverse microbial communities in humans, comprising more than 600 prevalent species (*25*). Distinct spatial structures, including “corn cob” structures, formed by spherical cells in consortium with a long filamentous core cell, and “hedgehog” structures have been previously discovered in the oral biofilm. The formation of these spatial structures is thought to be primarily driven by environmental and biochemical gradients, and potentially by metabolic cross-feeding between member taxa (*3*). We first designed a HiPR-FISH panel consisting of 233 probes targeting 54 genera from the human oral plaque microbiome. We observed corn cob structures decorated with *Streptococcus*, in agreement with previous observations (Figs. 4A-B). In addition to previously reported members of oral biofilm communities, such as *Porphyromonas* and *Corynebacterium*, we also observed bacteria that have been reported to be associated with periodontal disease, such as *Filifactor* (*26*), in consortium with other members of the human oral microbiota, including *Corynebacterium* and *Faecalibacterium*, highlighting the advantages of a probe design that is not biased by species abundance and the imaging multiplexity enabled by HIPR-FISH (Fig. 4C). We observed canonical hedgehog structures formed by *Corynebacterium* and took advantage of the single cell segmentation to measure the orientation of each cell relative to the horizontal axis of the image (Fig. 4D). In the field of view displayed in Fig. 4B, the filamentous cells gradually orient themselves from horizontal to vertical in the lower-left to upper-left direction. To further benchmark our probe design pipeline and the versatility of HiPR-FISH, we designed two additional HiPR-FISH probe sets, with different encoding sequences, barcode assignments, and probe selection criteria (see methods), consisting of 390 and 319 probes at the genus level targeting 65 and 61 genera, respectively. In experiments with all three HiPR-FISH probe sets, we observed many clusters of cells from the genus of *Lautropia* (Fig. 4E). In each design, spectra of *Lautropia* cells are cleanly resolved, and are classified correctly to the assigned barcode. As a further internal control, we compared the sizes of *Lautropia* cells detected using each design. We found that the cell sizes were consistent across all three different HiPR-FISH probe sets (Fig. 4F).

**Figure 4.**
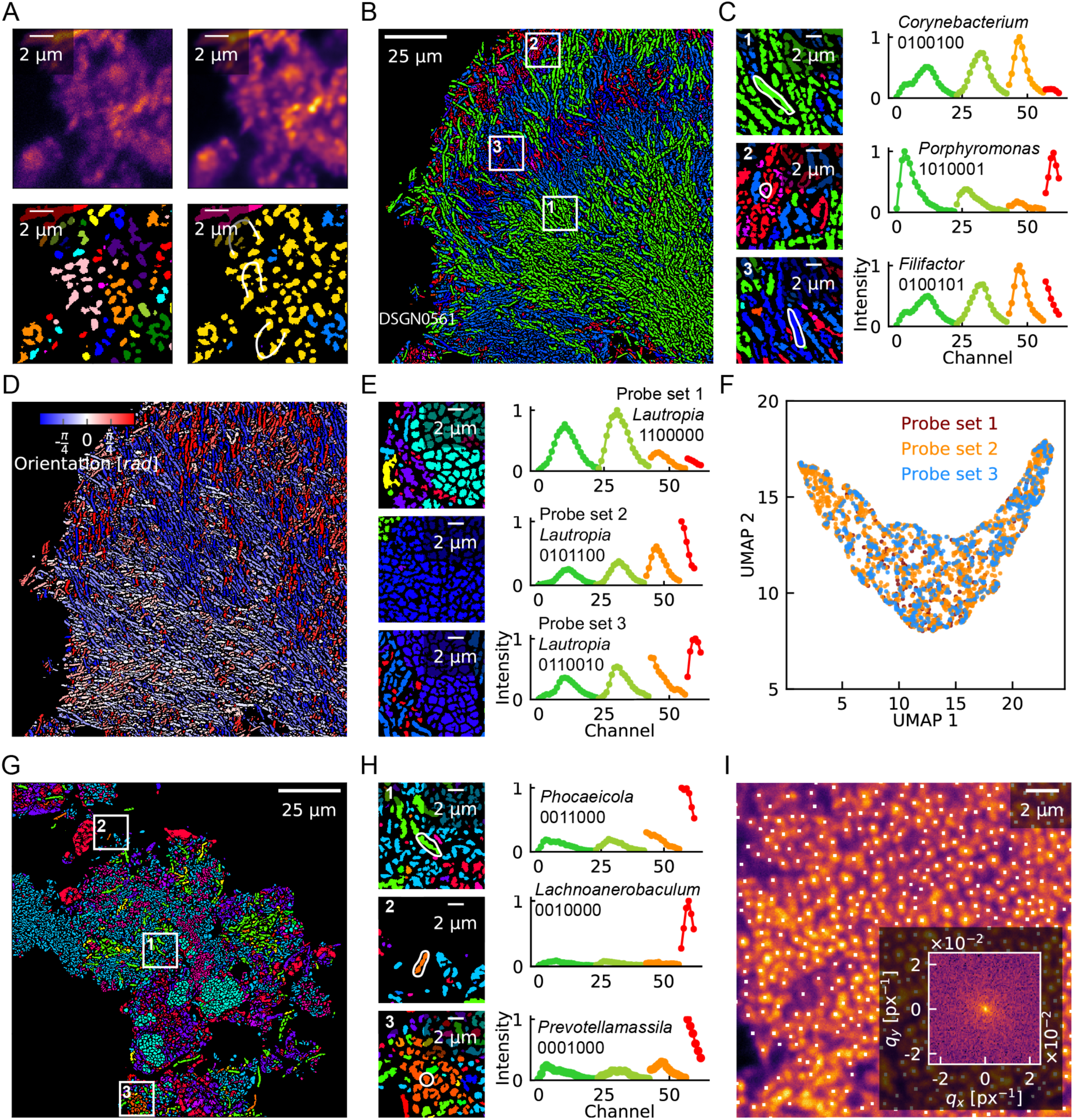
Biogeography of human oral biofilms. **A** Corn cob structures formed by cells from the genus of *Streptococcus* observed in human oral plaque biofilms. The raw image (upper left) was denoised (upper right), segmented (lower left), and identified (lower right). **B**. Corncob structures formed by cells from the genus of *Corynebacterium* (green cells) in human oral plaque biofilms. **C.** Magnified images of select areas in panel **B**, showing representative cells and their spectra. **D.** Single cell segmentation enables physical measurements, such as cell orientation. **E.** Example field of view and representative spectra of *Lautropia* cells observed using three probe sets with varying stringency. **F.** 2D UMAP projections of the physical properties of Lautropia cells observed using different probe designs. The physical properties used in the UMAP projection include semi-major axis length, semi-minor axis length, area, and eccentricity. Measurements from each probe panel are indistinguishable from each other. **G.** Cells from genera that were not targeted in previous studies were detected. **H.** Magnified images with representative cells and their spectra. **I.** Fourier transform of centroids of individual cells in clusters of *Actinomyces* cells shows signatures for an aperiodic structure with no short- or long-range order.

In addition to previously reported micro-architectures, we found structures not previously described. For example, we detected cells from the genus of *Phocaeicola* embedded among clusters of *Rothia* cells (Fig. 4G), suggesting the two genera interact metabolically in their native environment. We also observed cells from the genera of *Lachnoanerobaculum* and *Prevotellamassila* (Fig. 4H), which were not targeted by previous imaging experiments of the human oral microbiome, likely because of their low prevalence. The observed cell morphology for cells from the genus of *Lachnoanerobaculum* was consistent with those observed in bacterial culture based studies (*27*), with typical cell length ranging from 5 to 20 μm. Finally, we observed tight clusters formed by cells from the genus of *Actinomyces*. While the cell clusters appear to exhibit repetitive structures, Fourier transform of the image of single cell centroids suggests that the clusters are aperiodic, with no apparent short- or long-range order (Fig. 4I).

In the mouse gut microbiome, spatial relationships between the host and microbe as a whole have been reported. The biogeography of microbes in mock gut microbial communities have also been investigated, in experiments where germ-free mice were inoculated with a predefined consortium of microbes. The biogeography of the native mouse gut microbiome has however not been studied in its full complexity so far. Several studies have used shotgun and 16S rRNA sequencing to show that antibiotic treatment tends to reduce the gut microbiome diversity (*28*), but the effect of antibiotic treatment on spatial organization in the mucosa-associated gut microbial community has not been studied. To address these knowledge gap, we set out to create HiPR-FISH maps of the mouse gut microbiome for mice treated with two different antibiotics and controls. We designed two HiPR-FISH probe sets consisting of 115 and 264 probes targeting up to 47 genera. In the colon section from a mouse treated with Clindamycin, we detected cells from the well-studied genus of *Bacteroides*, as well as cells from the recently discovered genera of *Macellibacteroides* and *Longibaculum* (Figs. 5A-B). Using the single cell segmentation, we calculated the pair correlation function (PCF) for cells from the genera of *Bacteroides* and *Hespellia*, representing Bacteroidetes and Firmicutes, two of the well-studied phyla in the gut microbiome (Fig. 5C). The PCF describes how the density of objects varies as a function of radial distance from a reference object, and is often used in statistical mechanics to characterize structural features in systems of particles (*29*). We observed a slow decay in the PCF for *Bacteroides* indicating that *Bacteroides* cells tend to form clusters at short distances (∽ 3.3 μm). In contrast, the PCF of *Hespellia* cells were more consistent with a random distribution. We next measured the distribution of single-cell distances from the mucosal boundary with these three genera and found that *Bacteroides* are enriched near the boundary, while *Hespellia* are distributed relatively uniformly (Fig. 5D). The distribution of *Bacteroides* cells observed here is consistent with order-level measurements reported earlier in healthy mice (*10*). Applying HiPR-FISH to mouse gut tissue, we observed differences in microbiome spatial organization in mice treated with different antibiotics (Fig. 5E). Single-cell segmentation enabled us to calculate a region adjacency matrix, and subsequently a spatial association matrix, where each matrix element corresponds to the number of contacts between cells from any given two taxa. Adapting concepts from differential gene expression analysis in transcriptomics, we calculated differential spatial association matrices between mice treated with and without antibiotics (Figs. 5F & G). Using this analysis, we found that antibiotics treatment is associated with specific alterations to the spatial organization in the mouse gut. For example, the spatial associations between *Turicibacter* and *Lactonifactor* were reduced in Clindamycin treated mice, while those between *Turicibacter* and *Morella* were enhanced in Ciprofloxacin treated mice. Together, these results show the potential importance of considering spatial biogeography, and not just species diversity and abundance, for microbiome analyses.

**Figure 5.**
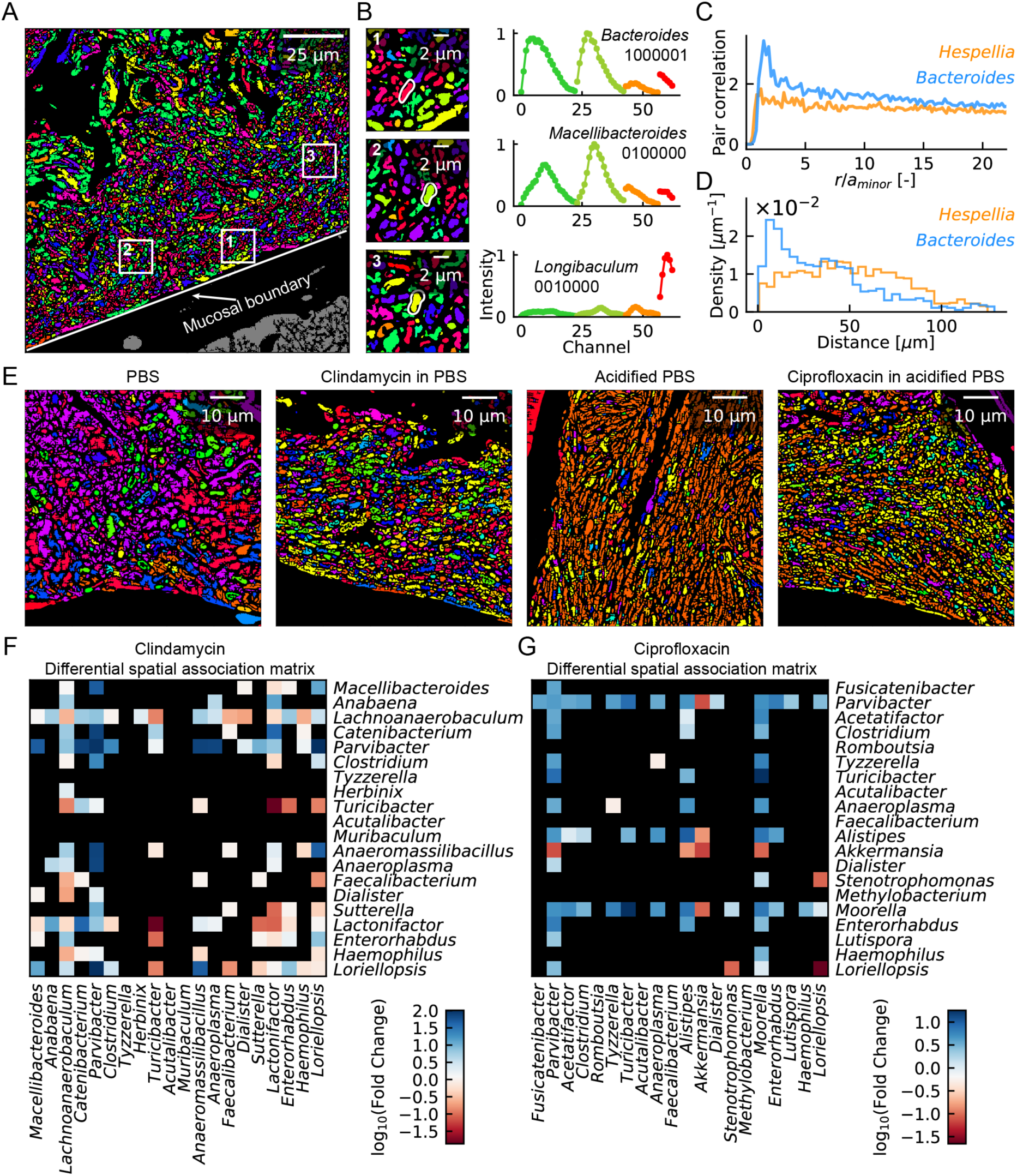
Biogeography of the mouse gut microbiome. **A.** A typical segmented and identified image of a mouse colon section. **B.** Magnified images of select areas in panel **A**, showing magnified views of representative cells and their spectra. **C.** Pair correlation function shows that cells in the genus of *Bacteroides* are likely to form clusters at short ranges, while *Hespellia* cells exhibit spatial distribution that are more consistent with a random distribution. The horizontal axis is normalized with respect to the average minor axis length of cells from each genus. **D.** Histogram of distances to the mucosal barrier reveals that *Bacteroides* cells are enriched near the mucosal boundary, while *Hespellia* cells are more evenly distributed away from the mucosal boundary. **E.** Example identified images of the mouse gut microbiome under different antibiotic treatments. **F & G.** Differential spatial association network analysis of the change in the number of contacts between cells from two taxa. Clindamycin treated mice shows reduced spatial association between *Turicibacter* and *Lactonifactor*, while ciprofloaxcin treated mice exhibits increased spatial association between *Turicibacter* and *Moorella*.

## DISCUSSION

Our experiments demonstrate that HiPR-FISH is a versatile tool to create spatial maps of microbial communities with high multiplexity. Compared to existing approaches, HiPR-FISH provides a more than tenfold improvement in the number of microbial taxa that can be tagged and identified (*11*) HiPR-FISH implements a two-step hybridization scheme that was previously exploited to spatially map messenger RNAs in tissues (*16, 30, 31*) In the context of microbiome mapping, this scheme greatly reduces the cost per assay design. Combined with the high abundance and relatively uniform distribution of 16S rRNA molecules in microbial cells, HiPR-FISH enables researchers to uniformly tag bacterial cells with combinations of up to ten fluorophores. HiPR-FISH requires only a single round of imaging on a commercial confocal microscope to decode multi-color barcodes *in-situ*. A single field of view can be imaged in just 5 minutes, which is much faster than FISH technologies that rely on multiple hybridization and imaging rounds (*16*). These experimental advantages make HiPR-FISH compatible with use cases that require fast data acquisition, such as applications in infectious disease diagnosis. For applications that do not rely on speed, implementations of HiPR-FISH with multiple rounds of hybridization and imaging can be considered, to further increase the multiplexity, to improve accuracy through the incorporation of error correction strategies (*16*), or to enable localization of bacterial transcripts or mobile genetic elements within HiPR-FISH maps.

HiPR-FISH provides a comprehensive framework for measuring microbial spatial organization. Measurements of the biogeography of complex microbial communities can provide deep insight into metabolic interactions between microbes and into interactions between microbes and their host. We expect that HiPR-FISH will have broad applicability in human health. HiPR-FISH can lead to entirely new avenues for investigations of complex microbial populations in the gut, in the oral cavity, or on implanted devices, all known to harbor biofilms. HiPR-FISH can also be applied to study gut-related disorders, such as Inflammatory Bowel Disease, where signaling between microbiota and gut epithelial tissue plays a role in reducing barrier integrity. HiPR-FISH can furthermore enable novel analyses of the role of the microbiota in the initiation and progression of tumors that form at epithelial barrier surfaces, such as colorectal cancer. Last, the quantitative single-cell measurements enabled by HiPR-FISH can become a rich resource for testing soft matter theories that describe physical principles governing microbial community assembly.

## METHODS

### Bacterial cell culture

All cultured cells were inoculated from frozen stock stored at −80°C into 4 mL of appropriate growth medium. A loopful of the liquid culture was streaked on an appropriate agar plate. A single colony from the agar plate was finally inoculated into appropriate liquid medium and grown to roughly mid-exponential phase. Details of growth media and growth time used are listed in Supplementary Table 1.

### Fluorescent readout probes

Fluorophore conjugated readout probes were purchased from Integrated DNA Technologies and Biosynthesis. Detailed information for each readout probe is listed in Supplementary Table 2.

### PacBio Sequencing

Metagenomic DNA from plaque samples were extracted using the QIAamp DNA Mini Kit according to manufacturer’s protocol. Ribosomal DNA was amplified from the extracted metagenomic DNA using the universal primers for the 16S rRNA listed in Supplementary Table 3, cleaned with the MinElute PCR purification Kit according to manufacturer’s protocol, and sequenced on a PacBio RSII or Sequel at the Duke Center for Genomic and Computational Biology. The PacBio sequence data was processed using rDnaTools (https://github.com/PacificBiosciences/rDnaTools), with a threshold of 99% accuracy. The output fasta sequences of the rDnaTools pipeline was used for probe design.

### Probe design and synthesis

The 16S rRNA sequences for cultured bacteria were generated using Sanger sequencing, while those for environmental samples were generated via PacBio sequencing. Probes were designed using a custom pipeline in python. All tools and packages used in the custom pipeline are listed in Supplementary Table 4. Briefly, the 16S sequences were grouped by taxon and sequence similarity. A consensus sequence was generated for each taxon using usearch. FISH probes for each consensus sequence were designed using primer3. The probes were then blasted against the database containing all 16S sequences from the community using blastn. A maximum continuous homology (MCH) score was calculated for each blast hit. The MCH score is defined as the maximum number of continuous bases that are shared between the query and the target sequence. Only blast hits above a threshold MCH score are considered significant and used for further analysis. The blast on-target rate and taxonomic coverage were calculated for each significant blast hit. Blast on-target rate is defined as the ratio between the number of correct blast hits and the total number of significant blast hits. The taxonomic coverage is defined as the ratio between the number of significant blast hits within the target species and the total number of sequences for the target species. Any probe with a blast on-target rate of less than 0.99 was excluded from the probe set to avoid ambiguity. For each taxon, the probe with the highest taxonomic coverage was then selected. Each probe was subsequently concatenated on both ends with 3bp spacers, readout sequences, and primer sequences. The 3bp-long spacers for each probe were taken from the three bases upstream and downstream from the target region on the 16S sequence of the probe. For each probe, all blast hits were examined for potential mis-hybridization sites. For each potential mis-hybridization site, we included a blocking probe that is complementary to the off-target sequence (*32*). Blocking probes were not conjugated to any readout sequences, and therefore did not contribute any fluorescent signals. The blocking probes were either purchased separately from Integrated DNA Technologies (www.idt.com) in plate format, or included in the complex oligo pool of encoding probes and purchased from CustomArrays (www.customarrays.com). The complex oligo pool was synthesized following the protocol as previously described (*16*). Briefly, the complex oligo pool was PCR amplified to incorporate T7 promoters, in vitro transcribed, reverse transcribed, and purified using ethanol precipitation.

### HiPR-FISH on synthetic *E. coli* communities

*E. coli* cells were grown overnight on LB agar plate. A single colony from the plate was inoculated into 800mL of LB broth supplemented with 40 mL of 1M potassium phosphate buffer and 40 mL of 20% glucose solution. Cells were grown for 7 hours to an OD of 1.1. Cultured cells were fixed for 1.5 hours by the addition of 800 mL of 2% freshly made formaldehyde. Fixed cells were aliquoted into 50mL tubes, concentrated by centrifugation at 4000 RPM for 15 minutes, resuspended in 1X PBS (1 mL per tube), and pooled together into two 50 mL tubes (∽16 mL of *E. coli* suspension each). Cell suspensions were washed 3 times in 1X PBS (50 mL per wash per tube), suspended in 50% ethanol, and stored at −20°C until use. Before the encoding hybridization experiment, cell suspension were treated with lysozyme solution (10 mg/mL in 10 mM Tris-HCl, pH 8) for 30 minutes at 37 °C, washed once with 1X PBS, and resuspended in 50% EtOH. Every 9.9 mL of encoding hybridization buffer includes 4 mL of cell suspension resuspended in 5.8 mL of Ultrapure water, 1 mL of 20X SSC, 1 mL of Denhardt’s solution, 2 mL of ethylene carbonate, and 100 µL of 1% SDS. Encoding hybridization buffers were aliquoted into 1.5 mL Eppendorf tubes at 99 µL per tube. Finally, 1 µL of the encoding probe for a barcode was added to each tube. The encoding hybridization suspension was briefly vortexed, incubated at 46°C for 4 hours, washed once in Washing Buffer (215 mM NaCl, 20 mM Tris-HCl, 20 mM EDTA) for 15 minutes at 48°C, washed twice in PBS, and resuspended in 100 µL of 50% EtOH. The 1023-plex synthetic community of barcoded *E. coli* was generated by mixing together 1 µL of each barcoded *E. coli* stock. To generate the titration community, we randomly divided the 1023 barcodes into 8 groups (7 communities with 128 barcodes each, and one community with 127 barcodes). Each 128-plex or 127-plex titration community was generated by mixing together a variable amount of barcoded *E. coli* stock. All synthetic community mixes were resuspended in 100 µL of 50% EtOH.

### HiPR-FISH on synthetic multi-species microbial communities

Cultured cells were fixed by adding an equal volume of 2% freshly made formaldehyde to the liquid culture for 1.5 hours. Fixed cells were washed 3 times in 1X PBS, permeated in absolute ethanol for 15 minutes, suspended in 50% ethanol, and stored at −20°C until use. For each species control experiment, 1 µl of 1 to 10 diluted pure culture was deposited onto a UltraStick slide and air dried. To permeate cell walls, we deposited 20 µL of 10 mg/ml lysozyme suspended in 10 mM Tris-HCl onto the slide and incubated it at 37°C for 3 hours. After lysozyme digestion, the slides were washed in 1X PBS for 15 minutes, dipped in pure ethanol, briefly rinsed with pure ethanol to remove any residual PBS, and air dried. The encoding hybridizations were performed in a 9 × 9 mm Frame-Seal hybridization chamber with 18 µl encoding hybridization buffer per slide at 46°C for 6 hours. The slides were then washed in the Washing Buffer at 48°C for 15 minutes, dipped in room temperature pure ethanol, rinsed with pure ethanol, and air dried. Readout hybridization were carried out at 46°C for 1 hour. The slides were washed and dried as described above and embedded in 15-30 µl Prolong Gold Anti-fade embedding medium.

### HiPR-FISH on human oral biofilm samples

Human plaque biofilm samples were obtained from a volunteer who refrained from oral hygiene for 24 – 48 hours. Subgingival plaque was collected using a stainless-steel dental pick, gently deposited into 1 mL of 50% ethanol, and stored at 4°C until use. For each human plaque experiment, 20 µL of plaque material was deposited onto an UltraStick slide and air dried. The slides were then fixed in 2% freshly made formaldehyde for 1.5 hours or overnight, washed in 1X PBS for 15 minutes, dipped in pure ethanol, rinsed with pure ethanol, and air dried. Lysozyme digestion,encoding, and readout hybridizations are carried out as described above.

### HiPR-FISH on mouse tissue

Mice were ordered from Jackson Laboratories and co-housed for 14 days. Mice were sacrificed using CO_2_ asphyxiation. The entire digestive tracks posterior to the stomach were fixed in Carnoy’s solution (60% ethanol, 30% chloroform, and 10% glacial acetic acid) for 48 hours at room temperature. Fixed tissues were rinsed three times in ethanol and stored in 70% ethanol at −20°C until paraffin embedding and sectioning. Tissues were embedded in paraffin and sectioned to 5 µm thickness. For deparaffinization, tissue sections on glass slides were incubated at 60°C for 10 minutes, washed once in xylene substitute for 10 minutes, once in xylene substitute at room temperature for 10 minutes, once in ethanol at room temperature for 5 minutes, and air dried. To reduce autofluorescence, deparaffinized slides were washed with 1% sodium borohydride in 1X PBS on ice for 30 minutes, with buffer change every 10 minutes, followed by three washes in 1X PBS on ice for 5 minutes each. Slides were briefly dipped in ethanol and allowed to air dry. Lysozyme digestion, encoding, and readout hybridization are carried out as described above.

### Spectral imaging

Spectral images were recorded on an inverted Zeiss 880 confocal microscope equipped with a 32-anode spectral detector, a Plan-Apochromat 63X/1.40 Oil objective, and excitation lasers at 405 nm, 488 nm, 514 nm, 561 nm, and 633 nm. The image acquisition settings are listed in Supplementary Table 5, 6, and 7.

### Flat field correction

To correct for the non-uniformity of the optical system across the field of view, we imaged a uniformly fluorescent flat field correction slide. One microliter of each readout probe was added to 90 µl of ProLong Glass embedding medium. The solution was vortexed and briefly centrifuged. Finally, 15 µL of the mixture was deposited onto a UltraStick slide, and a #1.5 coverslip was gently placed on top of the embedding medium. The flat field correction slide cured in the dark overnight and was imaged using the acquisition settings in Supplementary Table 5. Two fields of view of the flat field correction slide were averaged to generate the flat field correction image.

### Reference spectra measurement and classification

The reference spectrum for each barcode was measured using *E. coli* cells encoded with the corresponding barcode. For each barcode, *c*_*i*_ ≈ 300 – 500 single cell spectra in a single field of view were recorded. For each barcode, the average and standard deviation of the spectra was computed and used to simulate 5000 new spectra. Simulated spectra were used to train a classifier using a combination of support vector machine (SVM) and UMAP (Supplementary Fig. 1). In the first stage, four (7-bit experiment) or five (10-bit experiment) SVMs were trained to ascertain whether there was fluorescence signal in spectral images acquired using each of the lasers, which we refer to as channel signatures. The channel signatures were appended to the spectra as additional features to be used in UMAP projection. In the UMAP projection, we used a custom excitation-laser-wise cosine distance as a measure of distance. Given two spectra, we first compared the channel signatures. Spectra with different channel signature are assigned the maximum distance of 1. For spectra with the same channel signature, four or five (depending on how many excitation lasers were used) cosine distances were calculated, each corresponding to a cosine distance between spectra excited by the same laser. The final distance is the average of all channel-specific cosine distances. For synthetic community and environmental microbiome, reference spectra were simulated using only the spectra of individual fluorophores, taking into account Förster resonance energy transfer (FRET) and effects due to focus drift. For each fluorophore pair, we calculated the Förster distance using the emission spectra of the donor fluorophore and excitation spectra of the receptor fluorophore. To simulate FRET effects between fluorophores, we calculated FRET efficiencies between each fluorophore pair using a random distance between 6 to 10 nm. To simulate effects due to focus drift and potential fluorescence quenching due to interactions between fluorophores and other molecular motifs in the sample, we included a random excitation-laser dependent quenching factor q(λ_exc_) (Supplementary Table 8). To simulate background due to off-focus fluorescence signal and potential mis-hybridization, we simulated the background signal as the full-strength signal multiplied by 1 - q(λ_exc_). For each barcode, we simulated 2000 reference spectra. The simulated spectra were then trained using the same training architecture as before.

### Cultured cell imaging

For each field of view, spectral images were collected at five excitation wavelengths, and concatenated to form 95-channel images. The pixel size was 70 nm × 70 nm to ensure sampling at the Nyquist frequency. Each field of view is 2000 pixel × 2000 pixel in size, corresponding to a physical size of 135 µm x 135 µm.

### Biofilm imaging

Biofilms were imaged using the same spectral setting as described above, except for the laser power. All laser powers were changed by a common factor as necessary, to compensate for the overall lower intensity of environmental microbiome samples. For biofilm experiments, we only used the 488 nm, 514 nm, 561 nm, and 633 nm lasers. The pixel size was 70 nm × 70 nm to ensure sampling at the Nyquist frequency. For *z*-stack images, the step size along the optical axis was 150 nm.

### Image processing for cultured cells

The spectral classification and image processing pipeline are detailed in Supplementary Figure 1 and 2, respectively. Briefly, images acquired with each excitation laser were concatenated, registered, and denoised using non-local means or a convolution neural network. Denoised images were segmented using the watershed algorithm. For each cell, an average spectrum was calculated and assigned to the corresponding barcode using the spectra classification scheme described above.

### Image processing for biofilm samples

Biofilm images were acquired with each excitation laser. For volumetric images, multiple volumes of the same field of view were acquired with short pixel dwell time at low signal-to-noise ratio to reduce stage-drift induced motion blur. Raw volumes were computationally aligned to generate an average volume with minimal stage drift artefacts and high signal-to-noise ratio. For each voxel in the aligned volume, the line profile was extracted in multiple directions passing through the voxel under consideration. The structuring elements for the line profiles were parameterized by the azimuthal angle ϕ and polar angle θ. For 2D images, the azimuthal angle is set to zero. Each line profile was rescaled to the range [0,1], and the quartile coefficient of variation was calculated for each voxel. To produce the preprocessed image, the neighbor profile image was voxel-wise multiplied with the (1-quartile coefficient of variation). To distinguish signal pixels from background voxels in the pre-processed image, *k*-means clustering with *k* = 2 was used. A binary opening function was applied to the image to remove any residual connections between neighboring objects that have small number of connecting voxels. Any objects that are less than 10 pixels in size were removed, primarily to remove spuriously segmented objects in the background, and binary filling functions were used to fill in any holes in the segmented objects. Finally, the objects in the resulting image were labeled and served as the seed image for the watershed algorithm. To generate a mask image for watershed, the natural log of the raw volume averaged along the spectral axis was computed and *k*-means clustering with *k* = 2 was implemented to distinguish cells from the background. The intensity image for the watershed algorithm was simply the raw voxel averaged along the spectral axis. Finally, watershed segmentation was implemented using the intensity image, seed image, and the mask image generated above.

### Sequencing data availability

All PacBio sequencing data will be deposited to NCBI.

### Microscopy data availability

All microscopy data will be deposited to Zenodo.

### Code availability

All scripts will be available on Github.

## Materials and Correspondence

All requests should be submitted to I.D.V. and H.S.

## AUTHOR CONTRIBUTIONS

H.S. and I.D.V. conceived of the study. H.S., W.Z., I.B., and I.D.V. contributed to the study design. H.S. performed the experiments and analyzed the data. H.S., W.Z., I.B., and I.D.V. wrote the manuscript.

## ACKNOWLEDGEMENTS

We thank Rebecca Williams and Johanna M. Dela Cruz for assistance with microscopy. We thank Joan Lenz for assistance with bacterial culture. We thank Angela Douglas, John McMullen, Tobias Doerr, Joseph Peters, and Mike Petassi for generously providing materials. We thank Qiaojuan Shi for assistance with mouse work. We thank Phillip S. Burnham, Alexandre P. Cheng, Benjamin Grodner, Ti-yen Lan, and Elena Michel for discussions and feedback. This work was supported by US National Institute of Health (NIH) grant 1DP2AI138242 to IDV.

## Supplemental Materials

**Supplementary Table 1.** Information for species used in the synthetic community experiment.

| Species | Gram Stain | Medium | Temp. | Broth | Plate | Broth |
| --- | --- | --- | --- | --- | --- | --- |
| *Escherichia coli* MG1655 | Negative | LB | 37C | Overnight | Overnight | 16 hrs. |
| *Acetobacter tropicalis* | Negative | MRS | 30C | Overnight | Overnight | 16 hrs. |
| *Acetobacter pomorum* | Negative | MRS | 30C | Overnight | Overnight | 16 hrs. |
| *Lactobacillus brevis* | Positive | MRS | 30C | Overnight | Overnight | 16 hrs. |
| *Lactobacillus pomorum* | Positive | MRS | 30C | Overnight | Overnight | 16 hrs. |
| *Vibrio albensis* | Negative | LB | 37C | Overnight | Overnight | 16 hrs. |
| *Corynebacterium glutamicum* | Positive | TGY | 30C | Overnight | Overnight | 16 hrs. |
| *Acinetobacter schindleri* | Negative | TGY | 37C | Overnight | Overnight | 16 hrs. |
| *Comamonas testosteroni* | Negative | LB | 30C | Overnight | Overnight | 16 hrs. |
| *Enterococcus gallinarum* | Positive | MRS | 37C | Overnight | Overnight | 16 hrs. |
| *Xanthomonas vasicola* | Negative | TGY | 30C | Overnight | Overnight | 16 hrs. |

**Supplementary Table 2.**
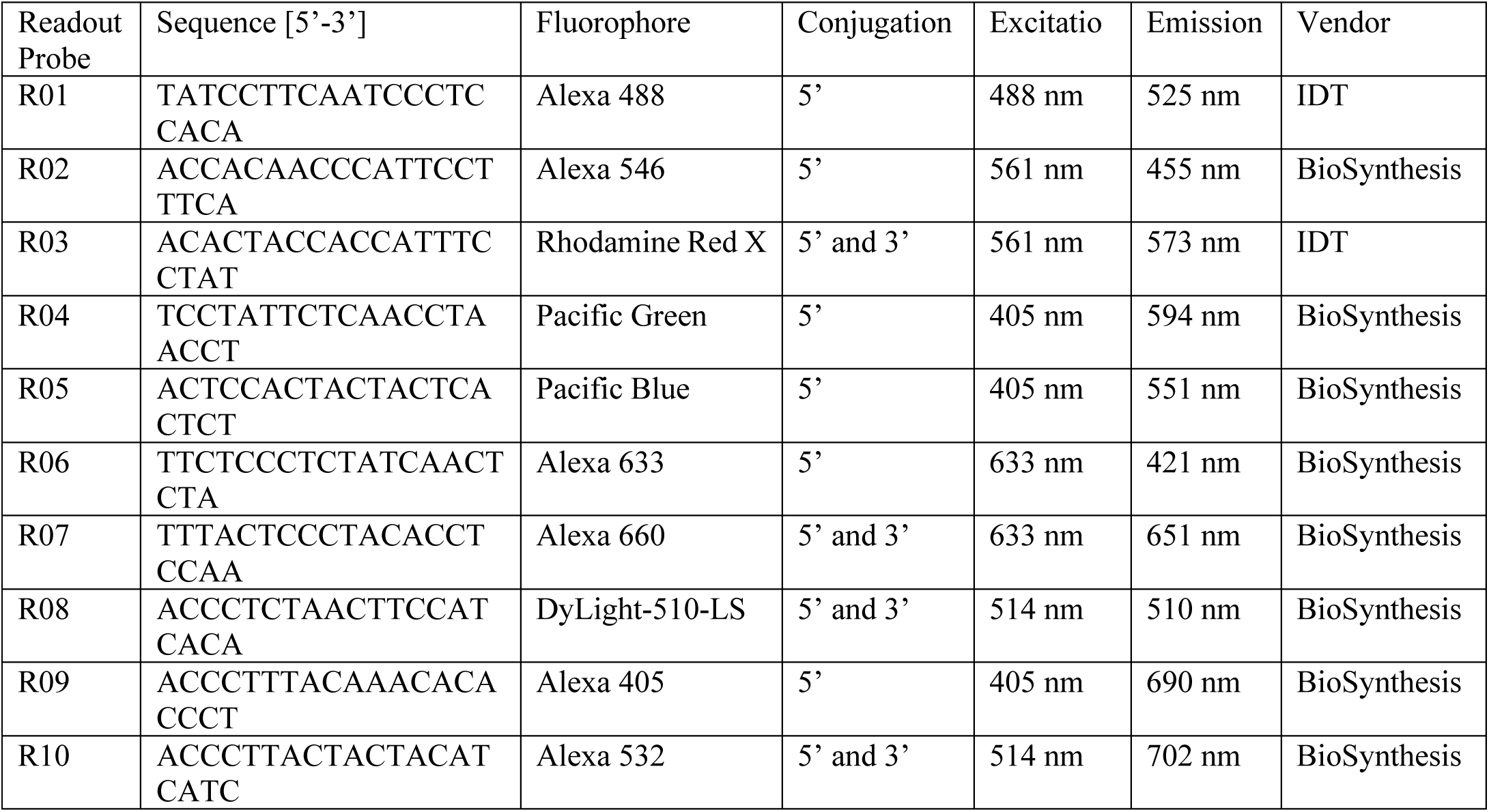
Information on readout probes.

**Supplementary Table 3.**
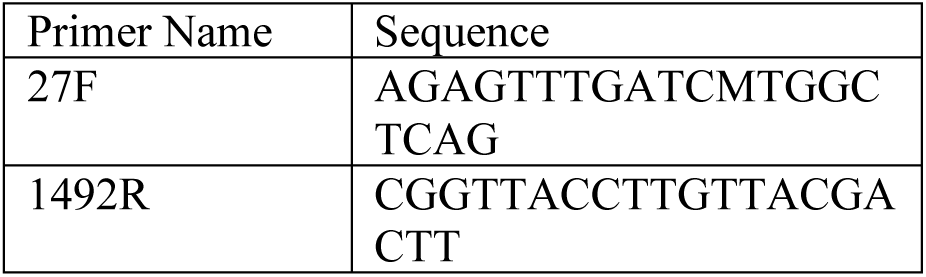
Universal primers for amplification of the 16S rDNA from metagenomic DNA.

**Supplementary Table 4.** Standalone software packages used in this work.

| Tool | Link |
| --- | --- |
| usearch | www.drive5.com/usearch |
| primer3 | https://sourceforge.net/projects/primer3/?source=directory |
| blastn | https://blast.ncbi.nlm.nih.gov/Blast.cgi?PAGE_TYPE=BlastDocs&DOC_TYPE=Download |

**Supplementary Table 5.** Microscope setting for 1023-plex *E. coli* experiment.

| Setting | Excitation | Laser Power | Master Gain | Emission Range | Emission Channel Width | Number of Emission Channels |
| --- | --- | --- | --- | --- | --- | --- |
| hiprfish_405 | 405 nm | 14 μW | 800 | 410nm – 695nm | 8.9 nm | 32 |
| hiprfish_488 | 488 nm | 11 μW | 800 | 491nm – 695nm | 8.9 nm | 23 |
| hiprfish_514 | 514 nm | 21 μW | 800 | 517nm – 695nm | 8.9 nm | 20 |
| hiprfish_561 | 561 nm | 11 μW | 800 | 517nm – 695nm | 8.9 nm | 14 |
| hiprfish_633 | 633 nm | 23 μW | 800 | 642nm – 695nm | 8.9 nm | 6 |

| Setting | Pixel Time | Scan Mode | Scan Direction | Beam Splitter | Averaging | Bit Depth |
| --- | --- | --- | --- | --- | --- | --- |
| hiprfish_405 | 2.11 μs | LineSequential | UniDirectional | MBS 405 | 4 | 8 |
| hiprfish_488 | 1.06 μs | LineSequential | UniDirectional | MBS 488 | 4 | 8 |
| hiprfish_514 | 1.06 μs | LineSequential | UniDirectional | MBS 458/514 | 4 | 8 |
| hiprfish_561 | 1.06 μs | LineSequential | UniDirectional | MBS 488/561 | 4 | 8 |
| hiprfish_633 | 1.06 μs | LineSequential | UniDirectional | MBS 488/561/633 | 4 | 8 |

**Supplementary Table 6.**
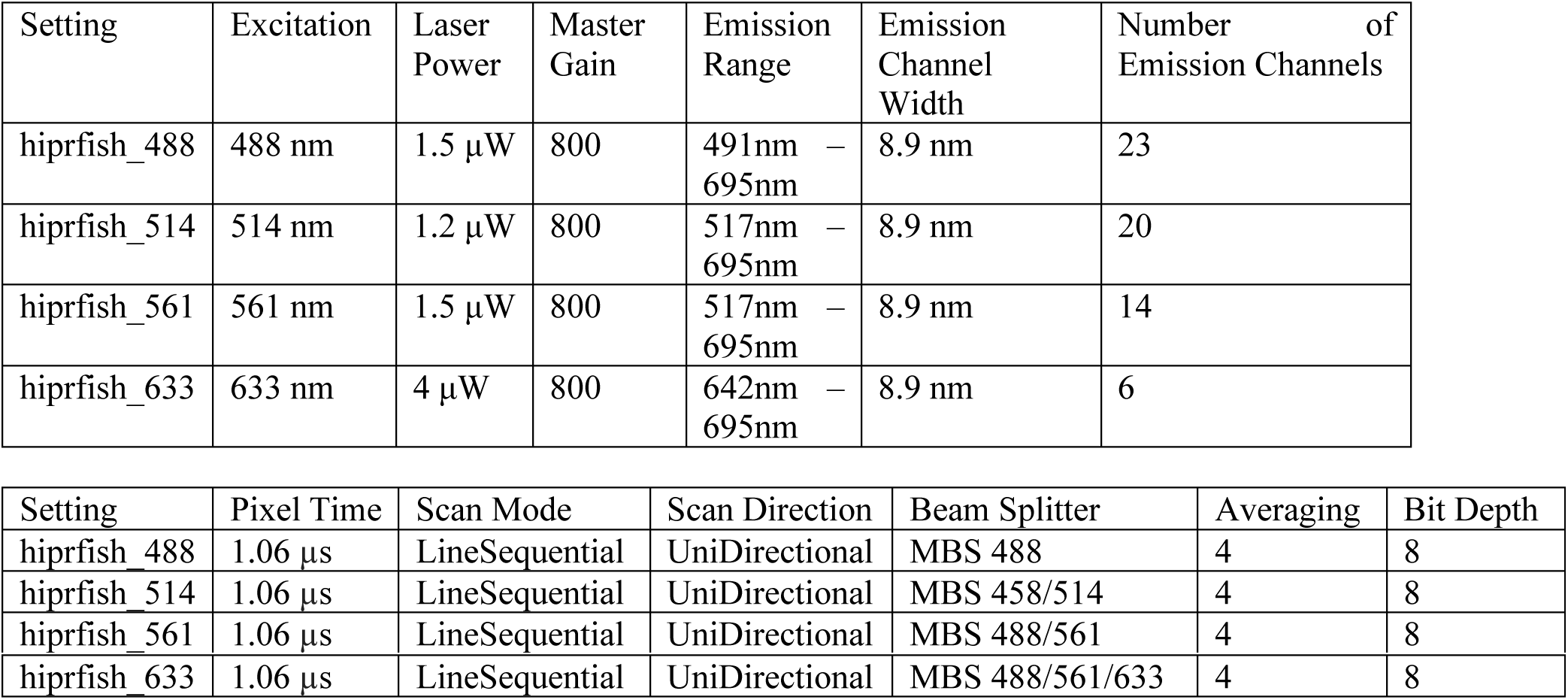
Microscope setting for 11-species synthetic community experiment.

**Supplementary Table 7.**
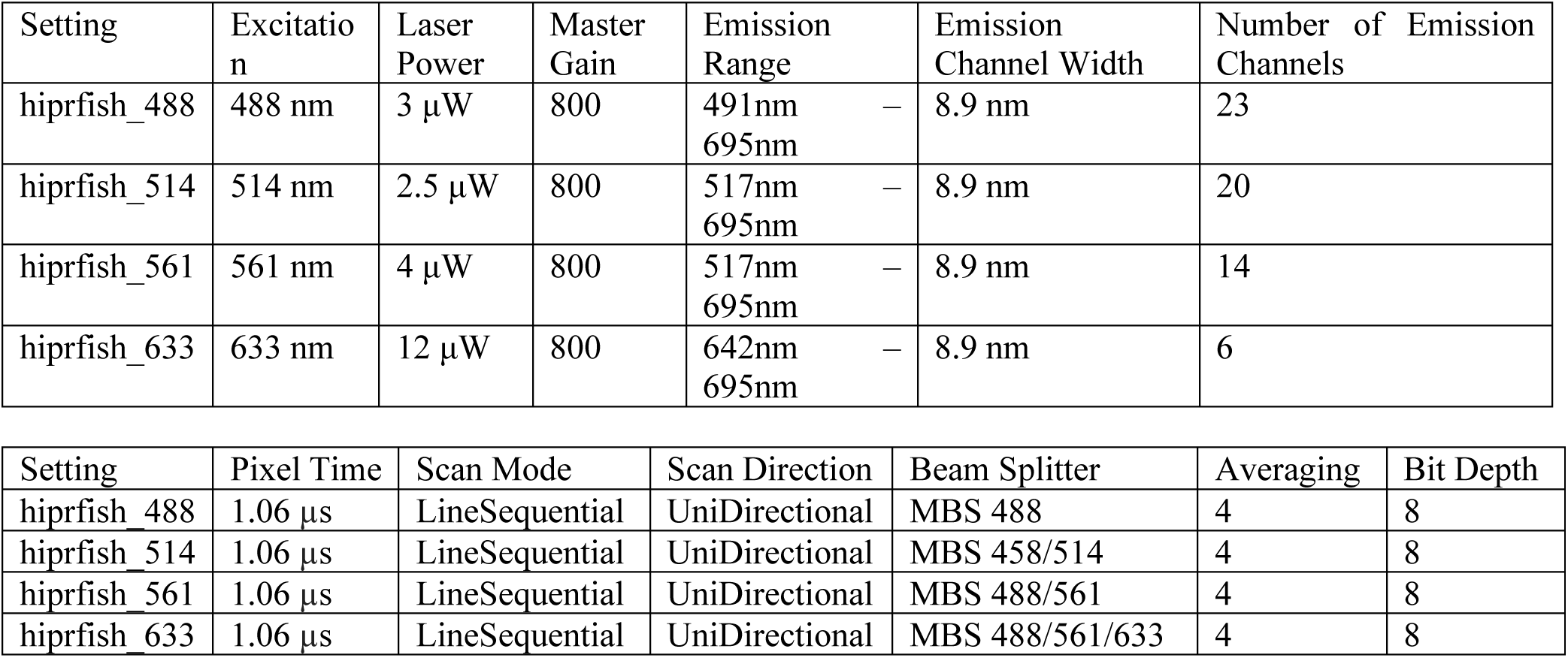
Microscope setting for environmental microbiome experiment.

| Setting | Excitation | Laser Power | Master Gain | Emission Range | Emission Channel Width | Number of Emission Channels |
| --- | --- | --- | --- | --- | --- | --- |
| hiprfish_488 | 488 nm | 3 μW | 800 | 491nm – 695nm | 8.9 nm | 23 |
| hiprfish_514 | 514 nm | 2.5 μW | 800 | 517nm – 695nm | 8.9 nm | 20 |
| hiprfish_561 | 561 nm | 4 μW | 800 | 517nm – 695nm | 8.9 nm | 14 |
| hiprfish_633 | 633 nm | 12 μW | 800 | 642nm – 695nm | 8.9 nm | 6 |

| Setting | Pixel Time | Scan Mode | Scan Direction | Beam Splitter | Averaging | Bit Depth |
| --- | --- | --- | --- | --- | --- | --- |
| hiprfish_488 | 1.06 μs | LineSequential | UniDirectional | MBS 488 | 4 | 8 |
| hiprfish_514 | 1.06 μs | LineSequential | UniDirectional | MBS 458/514 | 4 | 8 |
| hiprfish_561 | 1.06 μs | LineSequential | UniDirectional | MBS 488/561 | 4 | 8 |
| hiprfish_633 | 1.06 μs | LineSequential | UniDirectional | MBS 488/561/633 | 4 | 8 |

**Supplementary Table 8.** Excitation-laser dependent quenching factor q(λ_exc_).

|  | 488 nm | 514 nm | 561 nm | 633 nm |
| --- | --- | --- | --- | --- |
| R1 present | 0.25 | 0.25 | 0.35 | 0.45 |
| R1 absent | 0.05 | 0.25 | 0.35 | 0.45 |

**Supplementary Figure 1.**
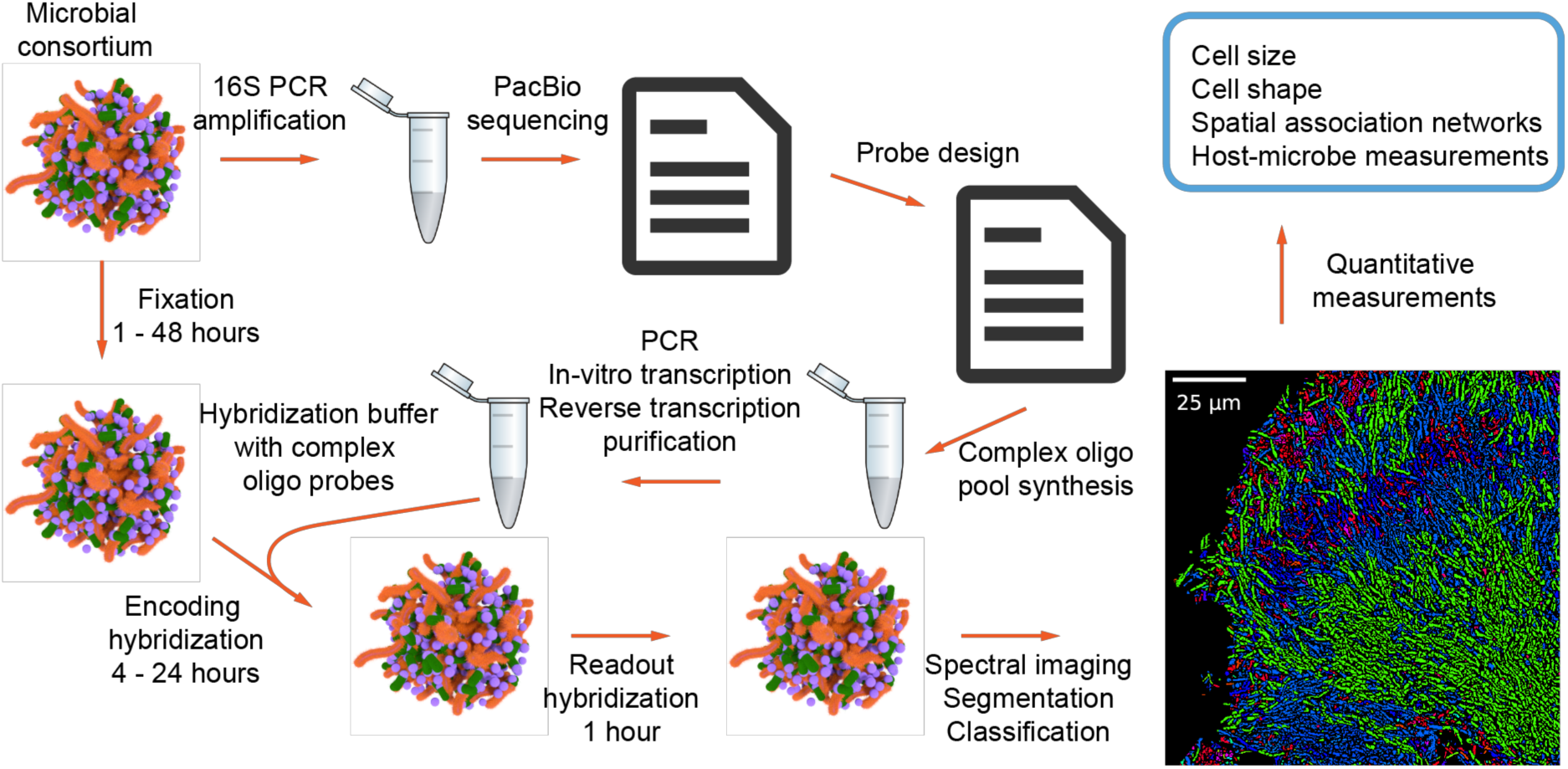
Typical workflow of HIPR-FISH. Environmental microbial consortiums are first split into two samples. One sample is used to generate full length 16S amplicon sequences using PCR and PacBio sequencing. The resulting sequence file is used to generate a list of probes, which are purchased from a commercial vendor. The other sample is used for imaging experiments. Fixed samples are hybridized using an encoding hybridization buffer containing the amplified complex probes and read out using a readout hybridization buffer containing fluorescently labeled readout probes. Samples are then embedded and imaged on a standard confocal microscope in the spectral imaging mode. Resulting raw images are registered and segmented. The spectra of individual cells are measured using the raw image and the segmentation and classified using a machine learning algorithm. Finally, classified images can be used for downstream quantitative measurements of microbial spatial associations.

**Supplementary Figure 2.**
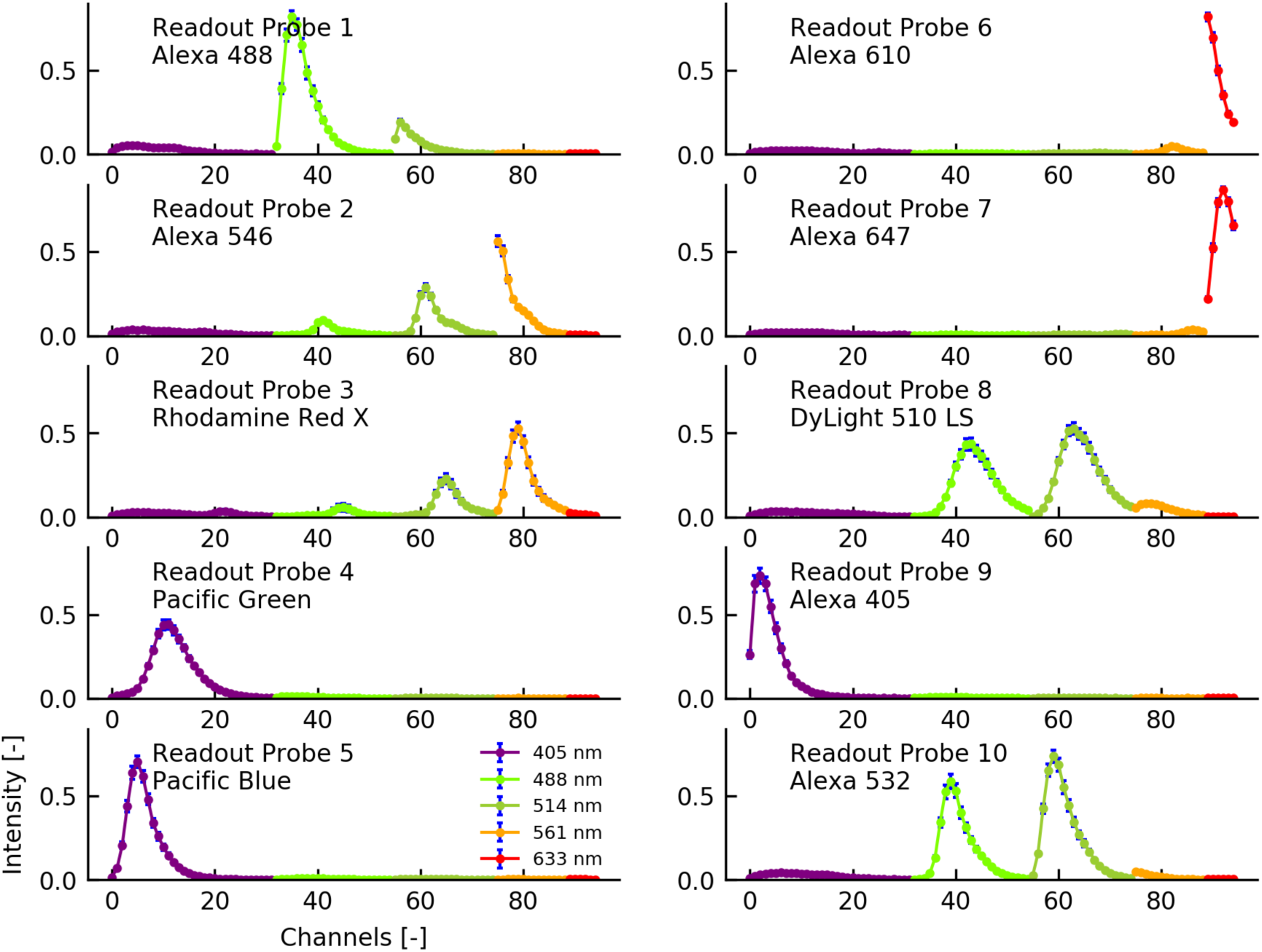
Spectra for the 10 fluorophores used in this study. The spectra were measured using *E. coli* cells labeled with Eub338 probes conjugated to each of the fluorophores. For each field of view, five excitation lasers are used to sequentially excite the fluorophores. The resulting spectral images are registered and segmented. Single cell spectra are measured using the raw images and the segmentations. Data points in this figure are intensity values averaged over all single cells in the imaged field of view. Error bars correspond to standard deviations of the measured single cell spectra.

**Supplementary Figure 3.**
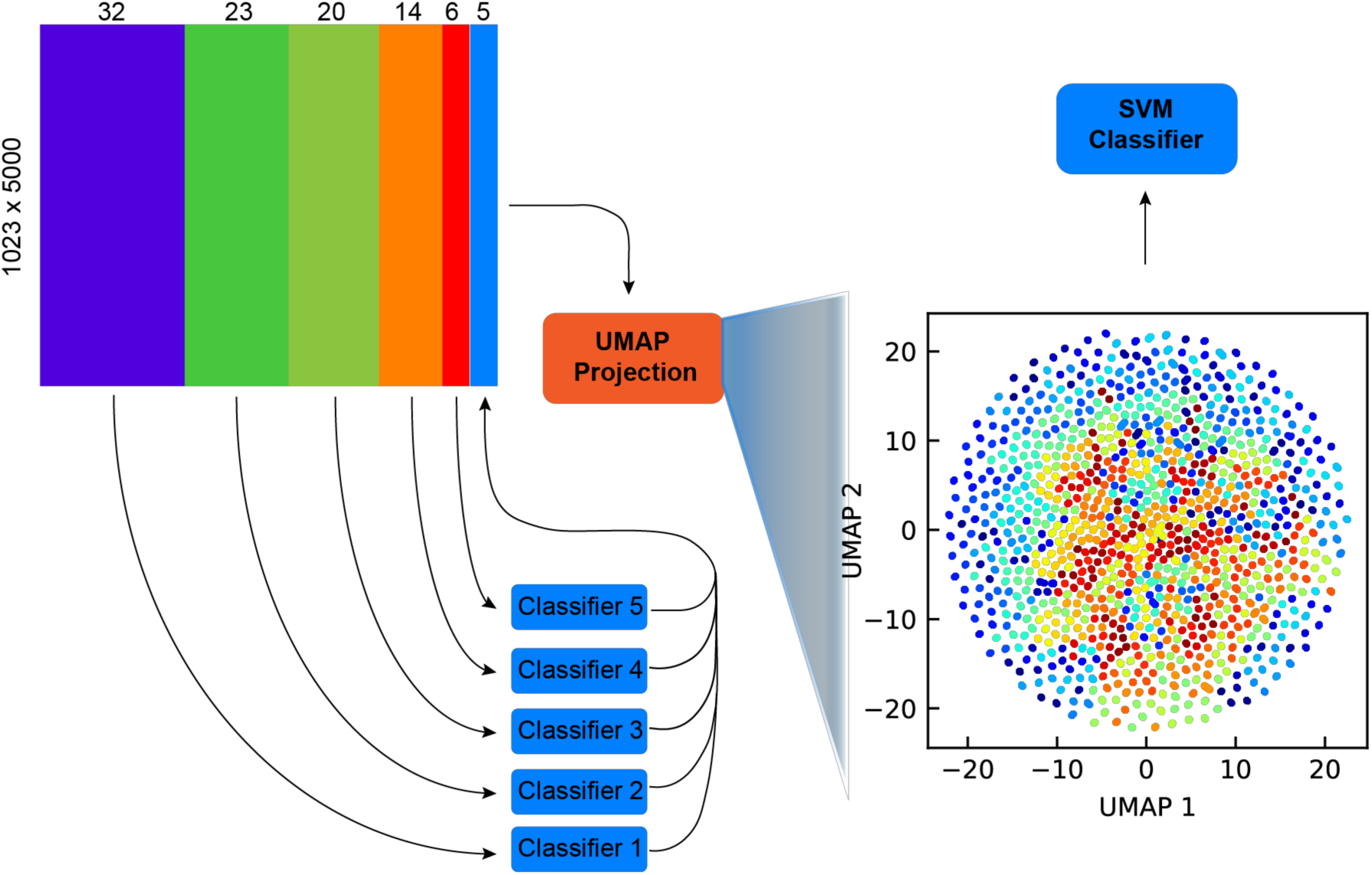
Algorithm for barcode classification using Uniform Manifold Approximation and Projection and a support vector machine classifier. For spectra acquired with each laser, we train a support vector machine to decide whether the detected spectra are signal or background. The five-column matrix indicating the presence or absence of signal in spectra acquired with each laser is concatenated with the spectra and sent to a UMAP transform with a custom defined metric. The custom defined metric is defined as the average cosine distance between two spectra acquired using the same laser. A final support vector machine is trained on the two-dimensional UMAP projection of the high-dimensional spectral data.

**Supplementary Figure 4.**
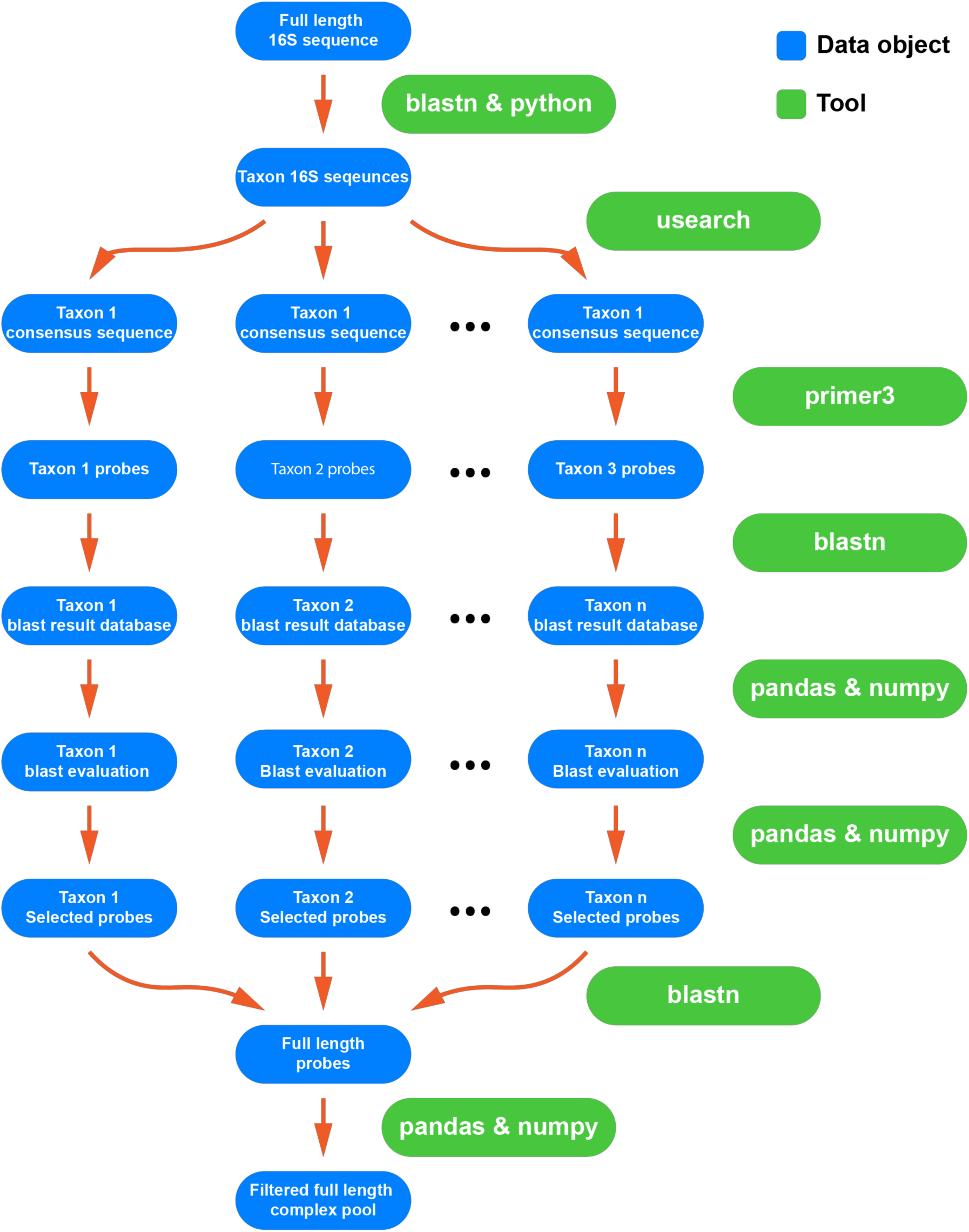
Schematic of the probe design pipeline. Full length 16S sequences are first grouped by taxa. The consensus sequence for each taxon is used to design probes. Each probe within each taxon is then blasted against the database of full length 16S sequences. Several probe quality metrics are calculated based on the blast results and are used to select probes. All selected probes are conjugated to the appropriate readout sequences, blasted against the database of full length 16S sequences to remove probes with any potential mis-hybridization sites due to the conjugation of the readout sequences.

**Supplementary Figure 5.**
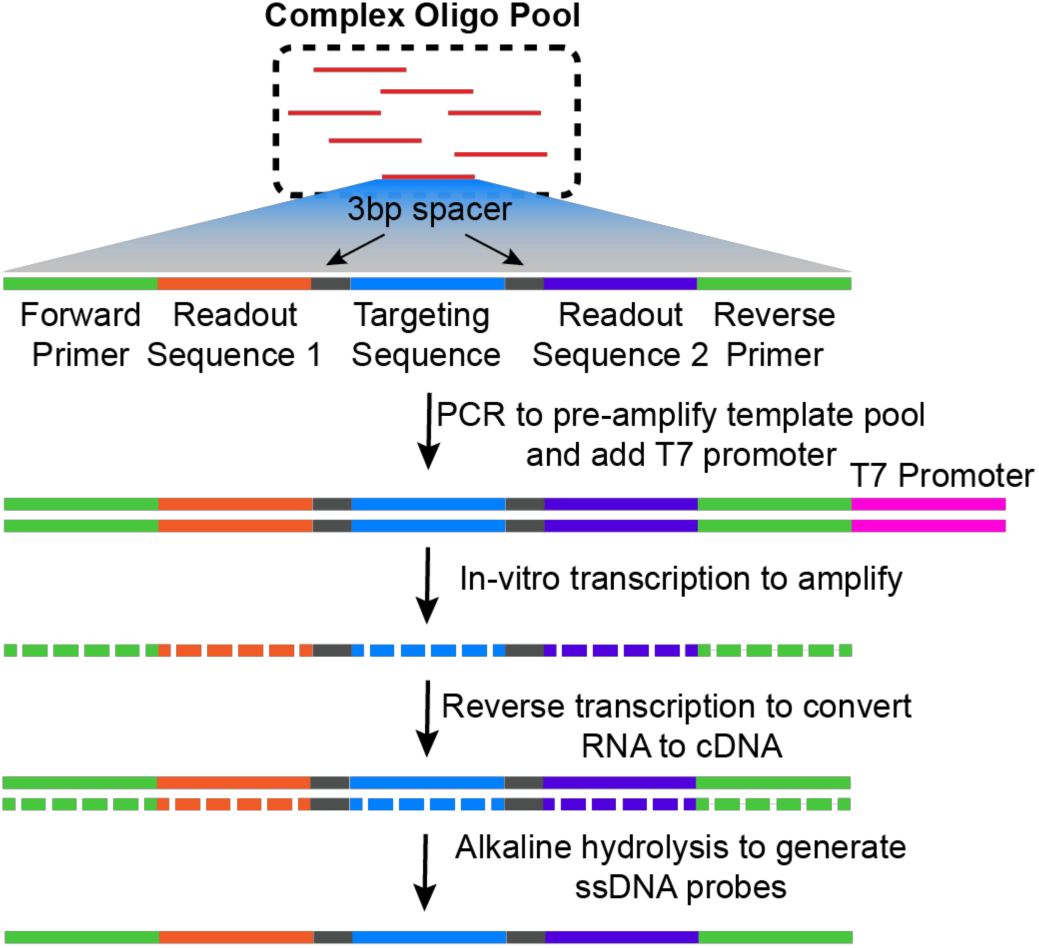
Schematic for probe synthesis. Complex oligo pools are amplified using limited cycle PCR. The T7 promoter introduced during PCR allows the templates to be in-vitro transcribed. Reverse transcription then converts RNA to cDNA. Finally, alkaline hydrolysis removes the RNA strand to generate the final single stranded DNA probe pool.

**Supplementary Figure 6.**
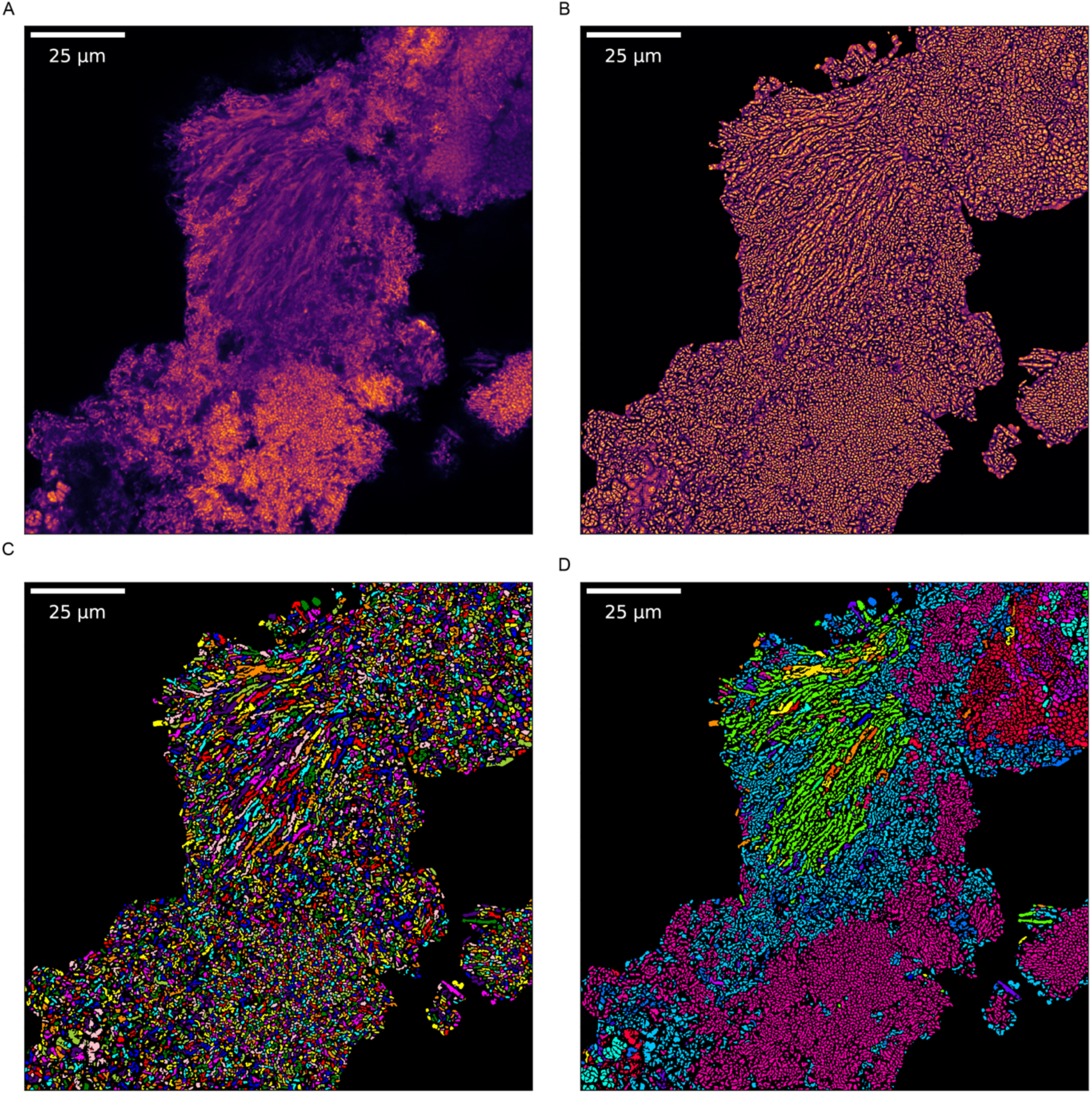
A. A typical raw image of a human plaque biofilm sample averaged along the spectral axis. **B.** The averaged image is then used to calculate an enhanced image using the local neighborhood line profile approach. **C-D.** The enhanced image enables watershed segmentation **(C)** and spectra classification **(D)**.

**Supplementary Figure 7.**
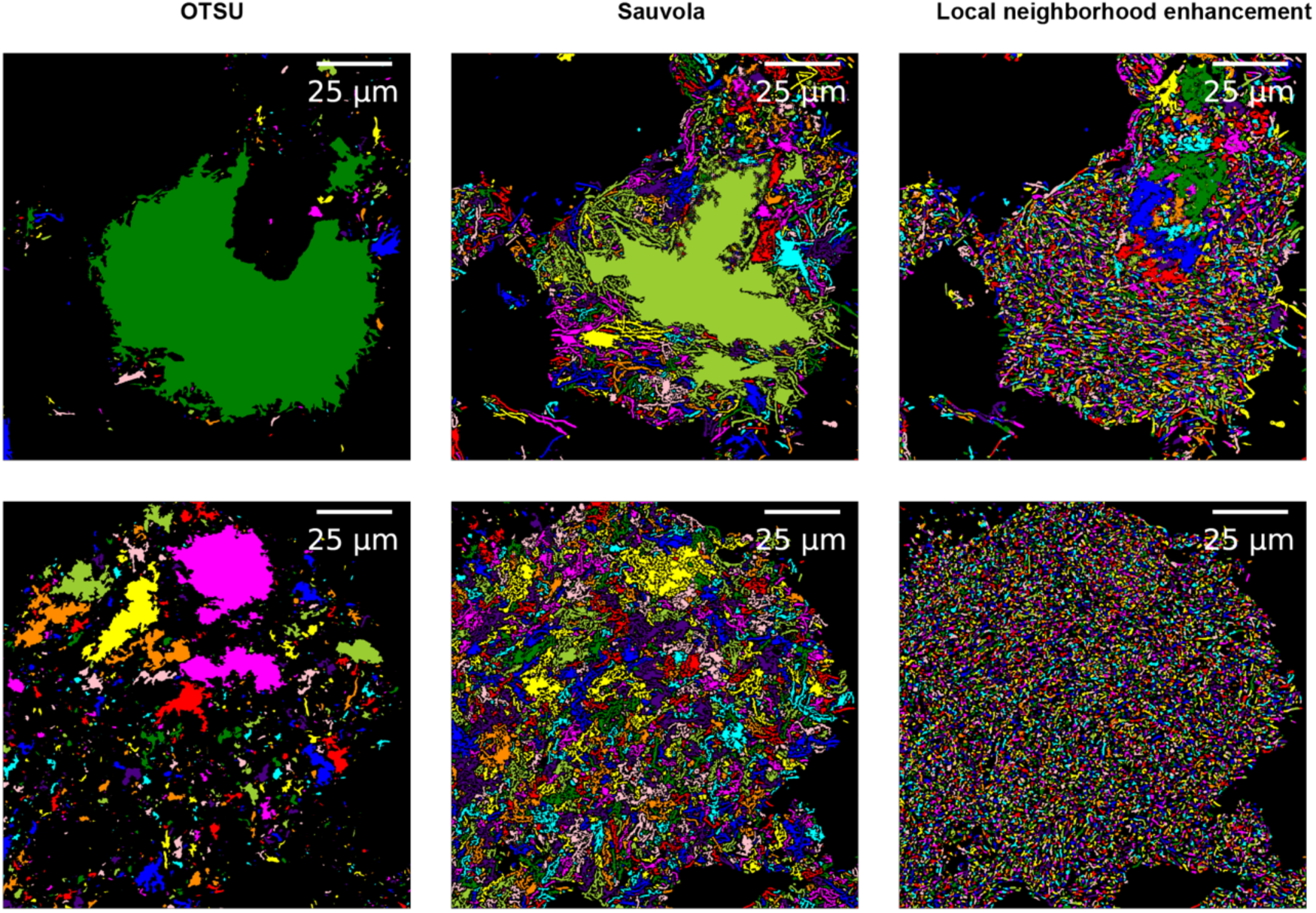
Comparison of approaches to segment microbial cells in biofilm images. Traditional adaptive thresholding techniques like Otsu’s thresholding performs poorly, while more advanced methods like Sauvola thresholding improve the accuracy of the segmentation. Our custom local neighborhood enhancement algorithm generates a much improved segmentation compared to prior approaches.

